# Unprecedented shift in Canadian High Arctic polar bear food web unsettles four millennia of stability

**DOI:** 10.1101/2022.11.28.518199

**Authors:** Jennifer Routledge, Christian Sonne, Robert J Letcher, Rune Dietz, Paul Szpak

## Abstract

Stable carbon (*δ*^13^C) and nitrogen (*δ*^15^N) isotope analysis was conducted on modern and archaeological polar bear bone collagen from the Canadian Arctic Archipelago to investigate potential changes in polar bear foraging ecology over four-millennia. Polar bear *δ*^13^C values showed a significant decline in the modern samples relative to all archaeological time-bins, indicating a disruption in the sources of production that support the food web, occurring after the Industrial Revolution. The trophic structure, indicated through *δ*^15^N, remained unaltered throughout all time periods. The lower *δ*^13^C observed in the modern samples indicates a change in the relative importance of pelagic (supported by open-water phytoplankton) over sympagic (supported by sea ice-associated algae) primary production. The consistency in polar bear *δ*^13^C through the late Holocene includes climatic shifts such as the Medieval Warm Period (MWP, A.D. 950-1250) and the early stages of the Little Ice Age (LIA, A.D. 1300-1850). These findings suggest that polar bears inhabit a food web that is more pelagic and less sympagic today than it was through the Late Holocene. We suggest that modern, anthropogenic warming has already affected food web structure in the Canadian Arctic Archipelago when modern data are contextualized with a deep time perspective.

**Research Highlights:**

1. Modern polar bear bone collagen *δ*^13^C suggests recent decline in ice associated prey.
2. Archaeological polar bear bone collagen suggests food web stability for millennia.
3. No change in sea-ice association through the Medieval Warm Period and Little Ice Age.
4. Recent isotopic shifts are unusual relative to the stability of ancient samples.
5. Modern Arctic warming has isotopically observable impacts that past events did not.

OR

**Significance Statement:** The lack of behavioral plasticity of both polar bears and their principal prey, ringed seals, make these species particularly vulnerable to declining sea ice. While the Lancaster Sound food web has demonstrated stability through past climate fluctuations, the speed and magnitude of ongoing changes in the Arctic has had an observable effect on the source of primary production. Given that past climate fluctuations are referenced as an argument to minimize the importance of modern anthropogenic warming, it is important to take opportunities to position contemporary climate change relative to the archaeological record. Here we present a unique illustration of the effects of past and present warming on polar bear diet and the marine food web in the Canadian Arctic Archipelago.

## Introduction

Polar bears (*Ursus maritimus*) are apex predators in a food web situated in a rapidly changing environment. Increasing global temperatures are causing precipitous circumpolar reduction in sea ice extent, with the magnitude of the effects in the Canadian Arctic Archipelago (CAA) increasing over the last two decades (Howell et al., 2016; IPCC, 2019; Noël et al., 2018;). Sea ice is an important feature of Arctic marine ecosystems, both as a physical habitat for marine mammals and as a substrate for primary production in the form of sea ice algae that bloom prior to pelagic phytoplankton (Leu et al., 2015). The landfast ice of the Canadian Arctic Archipelago hosts one of the most productive ice algae populations in the Arctic, driven by the input of nutrient rich waters from the Pacific Ocean (Leu et al., 2015).

In recent years, multi-year ice has given way to a thinner, seasonal ice regime and models of sea ice loss project the potential for ice-free Arctic summers by the end of the century, or likely sooner (Overland et al., 2019). Specialist, endemic predators like polar bears are expected to be among the first to face the impacts of a changing Arctic. Sea ice is required by polar bears for hunting their ice-obligate primary prey, ringed seals (*Pusa hispida*) (Laidre et al., 2008, 2022; Hamilton et al., 2017). When the sea ice breaks up in the summer, the bears enter a fasting period. The fasting period can be successfully endured for between 100 d for females with cubs, to more than 200 d for solitary males and females, provided they have adequate body mass at the onset of the fast (Molnár, 2020). Bears in several subpopulations are already approaching this fasting threshold (Molnár, 2020).

The ringed seal blubber and muscle consumed by polar bears during the spring hunt reflect the ice associated environment and compound specific stable isotope analysis (Kunisch et al., 2021) as well as a sea-ice specific isoprenoid lipid biomarker (IP_25_) (Brown et al., 2018) confirm the reliance of polar bear prey on the sympagic system. Through bulk stable isotope analysis, sources of primary production can be distinguished if their *δ*^13^C values are distinct. In the Arctic, sympagic (ice associated) algae have *δ*^13^C values 4-12 ‰ higher than pelagic (open-water) phytoplankton, making the two sources of primary production isotopically distinguishable (France et al., 1998; Søreide et al., 2006). There is little trophic enrichment of carbon isotopes in bone collagen, making δ^13^C a reliable indicator of the relative importance of primary producers (Deniro and Epstein, 1978; Fry and Sherr, 1989). Stable nitrogen isotope compositions (*δ*^15^N) undergo a predictable increase averaging approximately 3.4 ‰ with each trophic level (Post, 2002), providing a useful indicator of trophic structure.

Given the documented disruption to the Arctic cryosphere, it is reasonable to expect negative ice associated outcomes for polar bears. The Earth’s climate has never been static, however, and polar bears have endured past periods of warming and cooling. It is useful to contextualize the current crisis with the past to assess the magnitude of the impacts of anthropogenic warming relative to past climate disruptions (e.g., the Medieval Warm Period). Here, we compared the stable isotope compositions of modern and ancient polar bear bone collagen from the Lancaster Sound subpopulation (Figure 1) to assess the impacts on the polar bear’s food web ecology resulting from recent and past climate instability.

**Figure 1:**
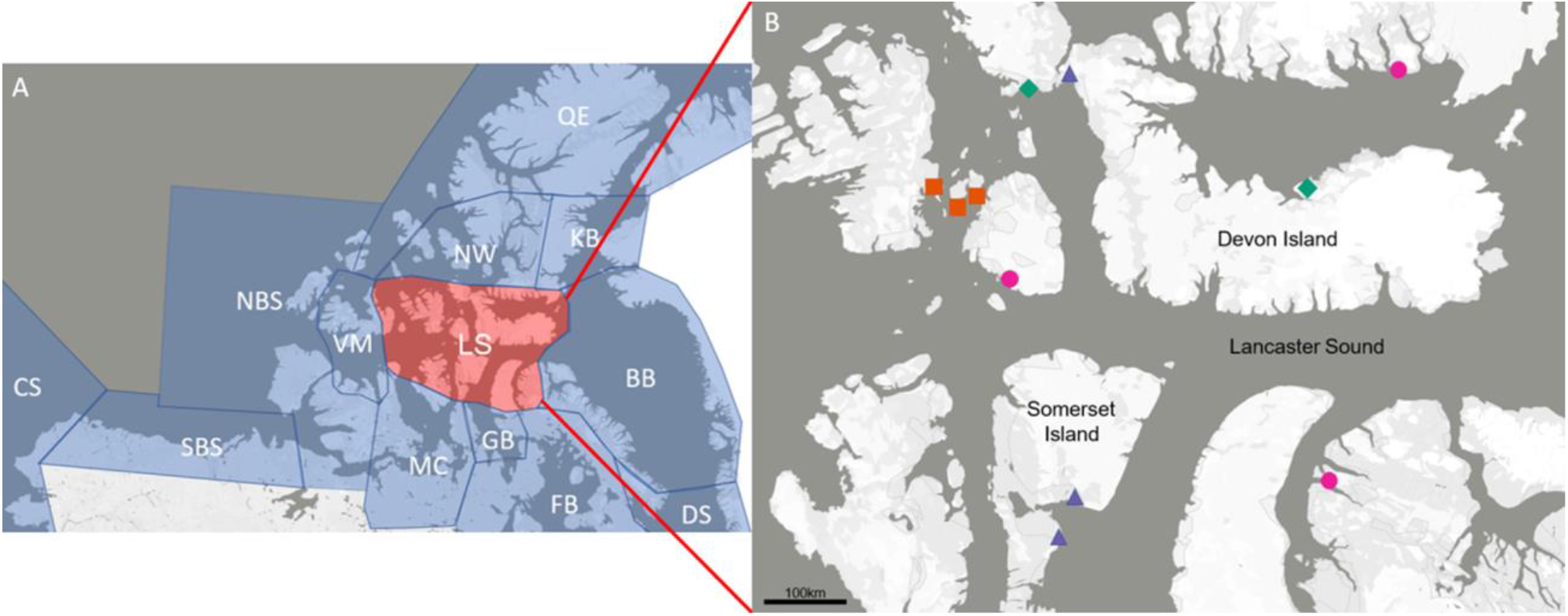
(A) Map showing the location of the Lancaster Sound (LS) polar bear subpopulation (red) relative to neighboring and nearby subpopulations (blue): Chuckchi Sea (CS), Southern Beaufort Sea (SBS), Northern Beaufort Sea (NBS), Viscount-Melville Sound (VM), M’Clintock Channel (MC), Gulf of Boothia (GB), Foxe Basin (FB), Davis Strait (DS), Baffin Bay (BB), Kane Basin (KB), Norwegian Bay (NW), Queen Elizabeth Islands (QE). (B) Map of the Lancaster Sound region showing the locations of the archaeological and modern sample acquisition sites: pre Dorset (green triangle), late Dorset (orange square), Thule (blue triangle), and modern (pink circle). Modern samples were collected within a 150 km radius of the communities indicated by the pink circles on the map.

## Results

The isotopic and elemental compositions of the modern and ancient polar bear samples are presented in full in the supplementary information (SI Table 2 and SI Table 3) and summarized in Table 1. Based on the atomic C:N ratios, the archaeological samples were assessed to be well preserved and uncontaminated (Deniro, 1985). The atomic C:N ratios of the modern samples fall within a very narrow range and after being corrected for lipid contamination comply with the most conservative quality control criteria established by Guiry and Szpak (2020) for modern bone collagen. The *δ*^13^C_corr_ of the modern samples was significantly lower than the *δ*^13^C of each of the ancient time periods (modern/Pre-Dorset *p* = 0.001, modern/Dorset *p* = 0.005, modern/Thule *p* = <0.001) (Table S1.1, Figure 2A). There were no significant differences in *δ*^13^C observed among any of the ancient time periods (Table S1.1). There was no significant difference in *δ*^15^N among any of the time periods (Table S1.1, Figure 2B). Polar bear bone collagen associated with the Dorset and Thule cultures preceding and including the MWP (respectively) did not produce significantly different isotopic compositions (Table S1.1, Table 1).

**Table 1.** Mean polar bear bone collagen *δ*^13^C and *δ*^15^N values for the four different time bins.

| **Time Period** | **Date Range** | **n Samples** | **$\delta^{13}\text{C}$**<br>**mean±1SD** | **$\delta^{15}\text{N}$**<br>**mean±1SD** |
| --- | --- | --- | --- | --- |
| Modern | 1998-2007 CE | 11 | $-13.93 \pm 0.17$ | $+21.86 \pm 0.68$ |
| Thule | 700-500 BP | 10 | $-13.30 \pm 0.39$ | $+22.03 \pm 0.73$ |
| Dorset | 1500-700 BP | 15 | $-13.56 \pm 0.41$ | $+22.28 \pm 1.29$ |
| Pre Dorset | 4000-2800 BP | 10 | $-13.36 \pm 0.39$ | $+22.48 \pm 0.99$ |

**Figure 2:**
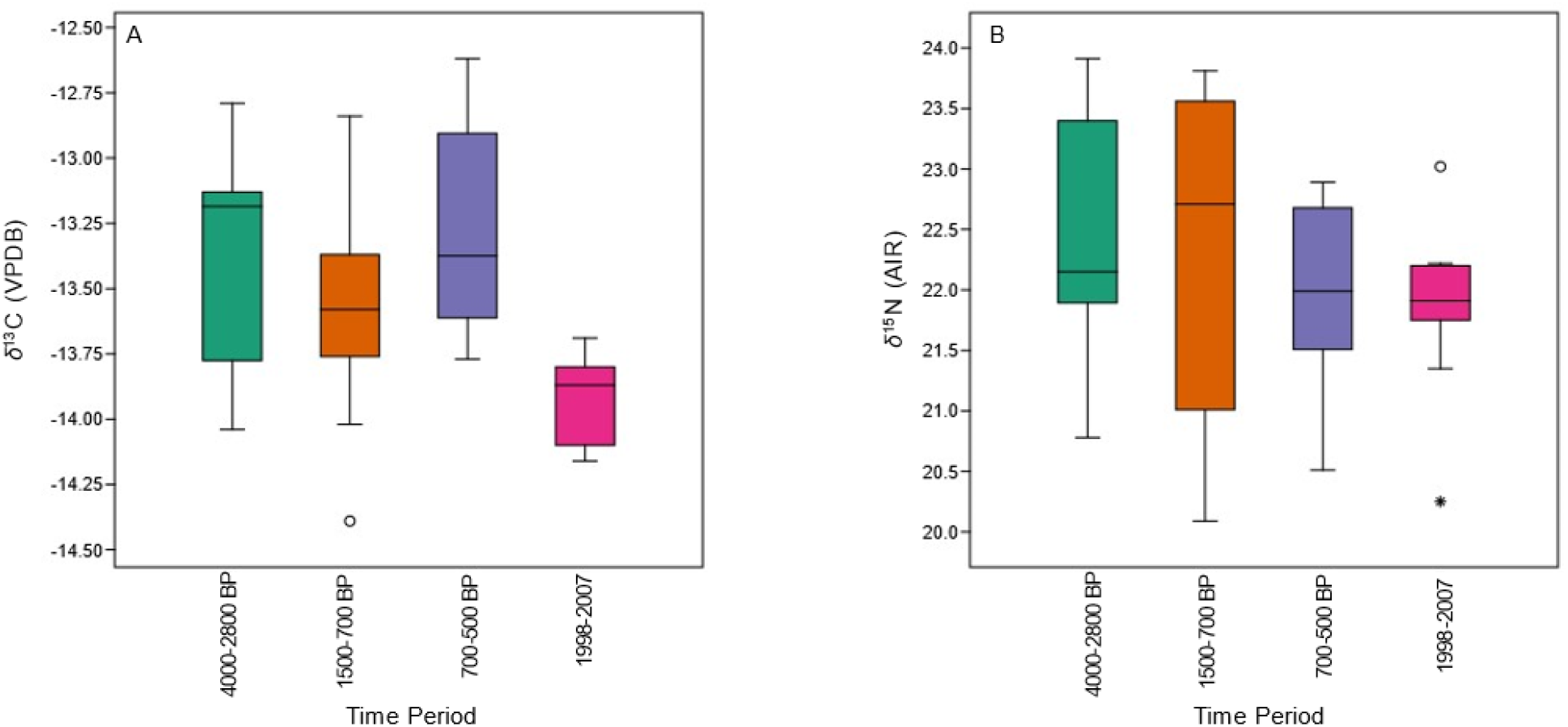
(A) Boxplot of polar bear bone collagen *δ*^13^C values. The modern *δ*^13^C values have been corrected for lipid contamination and the Suess Effect as described in the text. (B) Boxplot of polar bear bone collagen *δ*^15^N values.

## Discussion

The comparison of modern and archaeological polar bear bone collagen isotopic compositions indicates that there has been a recent shift in the Lancaster Sound food web, with modern polar bears presenting significantly lower *δ*^13^C values than ancient bears from any period. The lower *δ*^13^C values of the modern polar bears indicates a change in the source of primary production in the food web, specifically less exploitation of sympagic prey (with higher *δ*^13^C values) and a pivot toward greater consumption of pelagic prey. The difference persists after correction for the confounding factors of change to *δ*^13^C through CO_2_ produced by industrialization and endogenous lipid contamination. Both the CO_2_ produced by burning fossil fuels and lipids contaminating modern collagen shift the *δ*^13^C of modern samples lower and may be misinterpreted as more dramatic changes in polar bear diet or local ecological conditions without these adjustments. The differences observed here between modern and ancient samples are, therefore, conservative. The fact that a decrease in *δ*^13^C is still apparent for the modern bears, after accounting for these other factors, suggests a real environmental shift.

Similar observations have been made for modern and historic belugas in the Canadian High Arctic, with the interpretation of greater exploitation of the pelagic system (Desforges et al., 2021, Outridge et al., 2005), and in narwhal from northwest Greenland (Dietz et al., 2021). This interpretation is consistent with research on the quality and extent of sea ice in the Canadian Arctic Archipelago which indicates that seasonal consolidation in Lancaster Sound is increasingly incomplete, frequently consisting of mobile pack ice in open water throughout the winter (Haas and Howell, 2015). Assessment of the timing of autumn freeze-up and summer thaw indicated that from 1979 to 2014 the number of ice-free days has increased by an average of 1.08 d/yr (Regehr et al., 2016).

Under a scenario of increasing pelagic resource consumption, the magnitude of a shift in polar bear bone collagen *δ*^13^C should be relatively small since sympagic and pelagic primary producers often differ by only a few ‰ (France et al., 1998). A more pronounced decline in *δ*^13^C observed by McKinney et al. (2013) for East Greenland polar bear adipose sampled between 1985 and 2010 suggest an increasing reliance on pelagic prey over this short period (fewer ringed seals and more harp or hooded seals in the diet). This study, however, did not account for the Suess Effect so the magnitude of change may actually be more subtle or even non-existent. The fact that the isotopic data in our study were derived from bone collagen, which remodels at a very slow rate in mammals with its isotopic composition representing dietary intake over years (Hedges et al., 2007), means that short term temporal variation is more likely to have been dampened. The subtle temporal change that we observe in *δ*^13^C likely reflects real and sustained shifts in polar bear diets.

There was no significant change in *δ*^15^N over time indicating consistency in the trophic position of the bears through the Late Holocene and into the modern era. This is significant because it decreases the probability that polar bear *δ*^13^C values have declined due to a change in prey taxonomic composition. On the basis of quantitative fatty acid signature analysis, ringed seal, bearded seal, and beluga whale were interpreted to be the most important prey for Lancaster Sound polar bears (Thiemann et al., 2008). Relative to ringed seals, beluga whales in this region have comparable *δ*^15^N values but higher *δ*^13^C values (Szpak et al., 2019, 2020) while bearded seals have higher *δ*^13^C values and lower *δ*^15^N values relative to ringed seals (Jaouen et al., 2016). Neither species has the requisite lower *δ*^13^C values relative to ringed seals to explain the decline in polar bear *δ*^13^C in recent years (Figure 3). Moreover, the stability in polar bear *δ*^15^N across time is inconsistent with any appreciable variation in the importance of bearded seal in the diet since this would result in stronger temporal variation. The most likely explanation is a downward shift in the *δ*^13^C values throughout the food web caused by changes at the level of the abundance of sympagic and pelagic primary producers. While we do not have modern ringed seal *δ*^13^C data to speak directly to this hypothesis, Outridge et al. (2009) analyzed the *δ*^13^C values in ringed seal teeth from the 1300s, late 1800s, and 2002 and found the 2002 ringed seal *δ*^13^C values were ~2 ‰ lower than those from either the 1300s or late 1800s. Outridge et al. (2005) also found much lower (~4 ‰) *δ*^13^C values in beluga teeth from Somerset Island in the 1990s relative to the late 1800s. Neither of these studies extracted lipids from the samples, nor did they correct for the Suess Effect so the actual magnitude of changes in *δ*^13^C that they observed may be very similar to those observed here for polar bears. Nevertheless, the observation of a decline in tissue *δ*^13^C values for other high trophic level predators (including polar bear prey) in this environment support the hypothesis that a shift at the base of the food web is responsible for this pattern.

**Figure 3:**
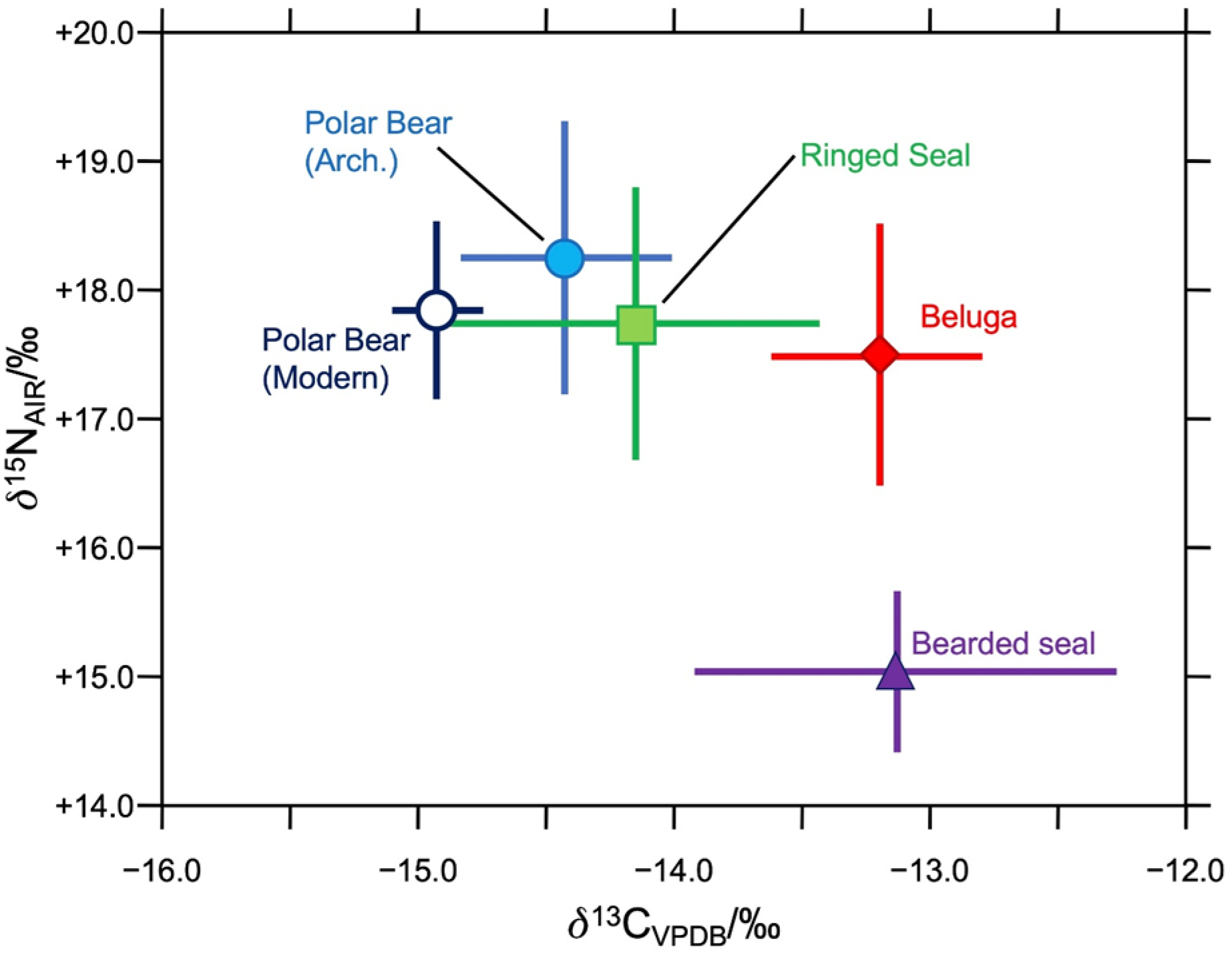
Stable isotopic compositions of modern (open circles) and ancient (filled circles) polar bears relative to isotopic compositions of the bone collagen of potential ancient prey. Bearded seal isotopic compositions (filled triangle) are derived from Late Dorset individuals (Jaouen et al., 2016). Beluga isotopic compositions (filled diamond) are from 19th century individuals from Somerset Island (Szpak et al., 2020). Ringed seal isotopic compositions (filled square) are from Thule individuals from Somerset Island (Szpak et al. 2019). Modern and ancient polar bear *δ*^13^C and *δ*^15^N values have been adjusted by –1.0 and –4 ‰, respectively, to account for the differences in bone collagen isotopic compositions between predator and prey (Bocherens and Drucker, 2003).

These previous studies were able to detect temporal shifts between two or three discrete points in time within the last 700 years. The polar bear data presented here extends this temporal framework four thousand years into the past. Notably, the modern polar bear *δ*^13^C values are lower than any of the preceding time bins. This implies that, while there may have been similar shifts in polar bear diets in the past, none of these seem to be of the magnitude that has been observed today. A very important caveat, however, stems from the derivation of the ancient samples from archaeological sites, meaning they represent bears that were hunted by humans, and were not evenly distributed across time. Based on extensive radiocarbon surveys (Savelle and Dyke, 2014; Savelle and Dyke, 2009; Savelle et al., 2012), large areas of the Canadian Arctic underwent massive demographic shifts with prolonged periods of abandonment. This is an important consideration because regional abandonments, potentially driven by unusual climatic conditions, would result in an absence of archaeological sites and consequently an absence of polar bear remains available to sample through these periods. We may, therefore, be missing important periods of climatic instability that were potentially unfavorable for both polar bear and human populations. Such a scenario is not inconceivable given that both polar bears and many of the human populations in the Canadian High Arctic relied to a great extent on ringed seals for subsistence (Darwent, 2001). Nevertheless, the stable carbon isotope compositions of Lancaster Sound polar bears from the 21st century are notably lower than any period for which we have data in the past four thousand years.

The significance of the modern shift in carbon isotopes can be illustrated by comparison within the ancient data set, between samples that precede the MWP (associated with Late Dorset archaeological sites) and samples dating to the MWP (associated with Classic Thule sites). If this past climate disruption was impactful to the food web, we would anticipate lower *δ*^13^C values during the MWP relative to the period immediately preceding it. We found no significant difference in the polar bear bone collagen *δ*^13^C values, suggesting relative stability in the importance of pelagic and sympagic prey immediately before and during the MWP. Decreased sea ice cover and the ability to pursue bowhead whales has been discussed as an important environmental driver of the expansion of the Classic Thule eastwards across the Canadian Arctic (McGhee, 1969/1970). Even if there were other non-environmental variables (e.g., demographic or social factors) wholly or partly driving the migration, other proxies for climatic conditions in the region suggest generally warmer conditions (Fisher et al., 1998; Thomas et al., 2010). Our data demonstrate that these relatively warm conditions did not result in a change in the isotopic composition of polar bear bone collagen and suggest that any changes in sea ice productivity during this time were subtle enough not to be reflected in the isotopic composition of polar bear tissues. The modern polar bear isotopic compositions are significantly lower than at any time in the past, suggesting an unusual environmental shift has taken place. The polar bear food web is derived in greater proportion from pelagic productivity now than at any past period for which an archaeological record exists in the region.

Due to a lack of foraging plasticity polar bears may be particularly vulnerable to declining sea ice extent. Ringed seals, the polar bear’s principal prey in the Lancaster Sound Region, are also impacted by declining sea ice as instability in their preferred denning habitat limits movement to refugia and impacts breeding success (Hamilton et al., 2016). While both of these high trophic level species have demonstrated resilience through past climate fluctuations, the speed and magnitude of change currently underway in the Arctic has already demonstrably impacted the food web as a whole and polar bears more specifically. Given a public discourse that occasionally invokes past climate fluctuations as an argument to minimize the importance of modern anthropogenic warming, it is important to take opportunities to position contemporary climate change relative to past events. By contextualizing the modern food web with observations of the same region over four millenia, the data presented in this study represent a unique illustration of the effects of past and present warming on the marine food web in the Canadian Arctic Archipelago.

## Materials and Methods

Ancient bone samples were collected from 35 polar bears from 10 archaeological sites (Supplement S2) in the region inhabited by the Lancaster Sound polar bear subpopulation (Figure 1). The archaeological samples came from pre Dorset (N= 10; 4000-2800 BP), Dorset (N= 15; 1500-700 BP), and Thule (N= 10; 700-500 BP) sites, and consisted of several different anatomical elements but each element represented a distinct individual (Table 1; Supplement S2). The archaeological samples could not be sexed or aged. Modern bone samples (N= 11; 1998-2007 CE) were obtained from 11 individuals harvested within a 150 km radius of the communities of Ikpiarjuk (Arctic Bay), Aujuittuq (Grise Fiord), and Qausuittuq (Resolute) between 1998 and 2007 (Supplement S3). The modern samples consisted of polar bear bacula, collected as a mandatory sample by subsistence hunters (Sonne et al., 2015). The modern samples were, therefore, male and ranged in age from three to 8-years-old (Sonne et al., 2015).

For the archaeological specimens, chunks of bone weighing ~200 mg were sampled using an NSK dental drill equipped with a diamond-tipped cutting wheel. Samples were demineralized in 16×100 mm glass culture tubes with 0.5 M HCl at room temperature. After the samples were demineralized, they were rinsed three times with Type I water (resistivity >18.2 MΩ·cm). The samples were treated with 0.1 M NaOH to remove humic contaminants from bones that exhibited a dark coloration. After 30 min, the NaOH was removed, and samples were rinsed twice with Type I water. The samples were then placed in a solution of 0.01 M HCl and placed in a dry bath at 75°C for 36 h, to solubilize the collagen. The collagen was extracted from the modern specimens using the same protocol with the following exceptions. First, powdered bone was removed from the samples using a Dremel equipped with a rotary burr. Because modern bone contains significant quantities of lipids whereas ancient bone does not, the samples were first treated with 2:1 chloroform:methanol under sonication for 1 h. The powdered samples were demineralized in 0.5 M HCl for 4 h. Demineralization in HCl does not alter the stable carbon or nitrogen isotopic composition of the bone collagen (Wilson and Szpak, 2022).

The solution containing the solubilized collagen was then transferred into pre-weighed 4 ml glass vials and freeze-dried. Collagen samples (0.45−0.55 mg) were weighed into tin capsules for analysis with a EuroEA 3000 (Euro Vector SpA) Elemental Analyzer coupled to a Nu Horizon (Nu Instruments) continuous flow isotope ratio mass spectrometer at the Water Quality Centre at Trent University. Ten percent of samples were analyzed in duplicate to assess homogeneity. Analytical sessions were calibrated using international standards USGS40, USGS41a and USGS66. Accuracy and precision were assessed with in-house check standards: SRM-1 (caribou bone collagen), SRM-2 (walrus bone collagen), and SRM-14 (polar bear bone collagen), the known isotopic compositions of which are given in supplement (S1). The standard uncertainty across analytical sessions was calculated to be ±0.13 ‰ for *δ*^13^C and ±0.24 ‰ for *δ*^15^N.

## Data Treatment

Modern bone samples may be particularly prone to contamination with lipids under certain circumstances (Guiry and Szpak, 2020). Despite lipid extraction with a commonly used protocol, we observed a correlation between the atomic C:N ratios and the *δ*^13^C values of the collagen even though our highest C:N ratio was 3.30 (Supplement S3). This observation is consistent with increasing quantities of residual lipids causing lower *δ*^13^C values as C:N ratios rise. No additional collagen from the modern specimens was available for further chemical pretreatment after the initial analyses. We therefore applied a mathematical correction to the modern bone collagen *δ*^13^C values (e.g., Post et al., 2007) using the following equation:

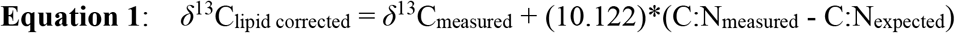

*δ*^13^C_measured_ and C:N_measured_ are the isotopic and elemental compositions determined for the sample prior to any adjustments. We used an expected C:N ratio for mammalian bone collagen of 3.23 (Guiry and Szpak, 2020) and 10.122 represents the slope of the regression line when plotting our modern C:N_measured_ and *δ*^13^C_measured_ (Supplement S1).

When comparing the *δ*^13^C values of modern and ancient samples, it is important to correct for the global decrease in *δ*^13^C of atmospheric and oceanic CO_2_ since the beginning of the industrial revolution (the Suess Effect; Eide et al, 2017; Francey et al., 1999). The magnitude of this effect on ocean surface water is lower than for the atmosphere and decreases with latitude (Sonnerup et al., 2000). A correction was calculated using Equation 2 (Hilton et al., 2006):

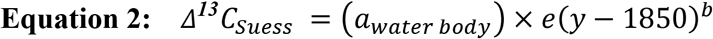

where α is the annual rate of decrease in *δ*^13^C specific to the water body (we used a value of 0.014 ‰ for the North West Atlantic (Mellon, 2018)), *y* is the year of sample harvest and *b* is the global oceanic decrease in *δ*^13^C (0.027 ‰; Gruber et al, 1999).

The carbon isotopic compositions of the modern samples have undergone correction through Equations 1 and 2 and the revised carbon isotope values are denoted *δ*^13^C_corr_. Both of these corrections have been applied to account for confounding causes of low *δ*^13^C values in modern samples to avoid an exaggeration of any differences between modern and ancient samples since both of these adjustments increase the *δ*^13^C values of the modern samples (Figure S1). Statistical comparisons were made using Past 4.03 (Hammer, 2020). Differences in polar bear isotopic compositions between periods were assessed using Welch’s t-tests after assessing the normality of the distributions with a Shapiro-Wilk test.

## Supporting information

Supplement S2 Archaeological Data

Supplement S3 Modern Data

