## Supplement S1 for "Unprecedented shift in Canadian High Arctic polar bear food web unsettles four millennia of stability"

**Supplementary Information for Routledge et al. Long Term Food Web Patterns in the Canadian High Arctic: Evidence from Stable Isotope Analysis of Polar Bears**

**Supplemental Materials and Methods**

**Summary of Statistical Analysis in this Study**

Differences of means for the four time-bins represented in this study were assessed through Past 4.03 (Hammer, 2020) using Welch's t-tests (Table S1.1).

Table S1.1 Results of Welch's t-tests comparing isotopic compositions for each time bin. Bold p-values indicate statistically significant differences.

| Variable | | Dorset | Thule | Modern |
|---|---|---|---|---|
| $\delta^{13}C$ | Pre-Dorset | 0.22 | 0.74 | **0.001** |
| | Dorset | | 0.12 | **0.005** |
| | Thule | | | **<0.001** |
| $\delta^{15}N$ | Pre-Dorset | 0.68 | 0.26 | 0.12 |
| | Dorset | | 0.53 | 0.29 |
| | Thule | | | 0.59 |

**Supplementary Materials and Methods**

*Addressing Potential Sample Bias*

The lack of diversity in sex (all male) and age (all adult) for the modern samples is a potential source of bias. Thiemann et al. (2008) studied polar bears across ten Canadian subpopulations and found that sex and age resulted in differential polar bear foraging in some but not all regions of the Arctic. In some cases, larger, older males had the most diverse diets relative to females and younger males. This pattern was not observed in Lancaster Sound, Baffin Bay, and Gulf of Boothia, where there was no indication of differential foraging by age. Additionally, bears from Lancaster Sound, Baffin Bay, and Davis Strait were not characterized by any differences in prey consumption according to sex. It is, therefore, unlikely that the biased sex distribution of our modern samples is the source of the differences in $\delta^{13}C$ between the ancient and modern populations but we cannot rule out the possibility that sex-based differences in diet may have existed in the past for the Lancaster Sound population that do not exist in today's population. In the future, sexing of the ancient polar bear remains via ancient DNA, combined with stable isotope analysis could be used to test this hypothesis (e.g., Szpak et al., 2020).

*Calibration and Analytical Uncertainty*

Elemental and isotopic compositions of carbon and nitrogen were obtained through analysis using a EuroEA 3000 (Euro Vector SpA) Elemental Analyzer coupled to a Nu Horizon (Nu Instruments) continuous flow isotope ratio mass spectrometer at the Water Quality Centre at Trent University. Measurements were calibrated relative to VPDB for $\delta^{13}$C and AIR for $\delta^{15}$N using USGS40, USGS41a and USGS66 (Table S1.2). Quality Assurance was conducted using internal check standards (Table S1.3) and sample duplication.

Table S1.2 Known Values of Calibration Standards

| Standard | Material | $\delta^{13}C_{VPDB}$/‰ | $\delta^{15}N_{AIR}$/‰ | Reference |
|---|---|---|---|---|
| USGS40 | Glutamic acid | −26.39 ± 0.04 | −4.52 ± 0.06 | Qi et al. (2003) |
| USGS44a | Glutamic acid | +36.55 ± 0.08 | +47.55 ± 0.15 | Qi et al. (2016) |
| USGS66 | Glycine | −0.67 ± 0.04 | +40.83 ± 0.06 | Schimmelmann et al., (2016) |

Table S1.3 Known Values of Internal Standards

| Standard | Material | N | $\delta^{13}C_{VPDB}$/‰ | $\delta^{15}N_{AIR}$/‰ |
|---|---|---|---|---|
| SRM-1 | Caribou bone collagen | 861 | −19.39 ± 0.08 | +1.83 ± 0.16 |
| SRM-2 | Walrus bone collagen | 442 | −14.81 ± 0.06 | +15.58 ± 0.15 |
| SRM-14 | Polar bear bone collagen | 776 | −13.64 ± 0.08 | +21.60 ± 0.24 |

*Data Treatment - Correction for Lipid Contamination*

The presence of a correlation between the atomic C:N ratios and the $\delta^{13}$C values of the modern polar bear bone collagen (Pearson's $r = -0.83$, $p = 0.002$) suggests that variable presence of lipid contamination (Figure S1). Mammalian collagen has a low atomic C:N ratio, around 3.23, while lipids contain primarily carbon with lower $\delta^{13}$C values than proteins from the same animal (DeNiro and Epstein, 1977) and little to no nitrogen. As the proportion of lipids in the sample increases, the $\delta^{13}$C value decreases and the C:N ratio increases; this relationship is fairly linear across a short range of atomic C:N ratios (e.g., 3.2-4.0; Guiry and Szpak, 2020).

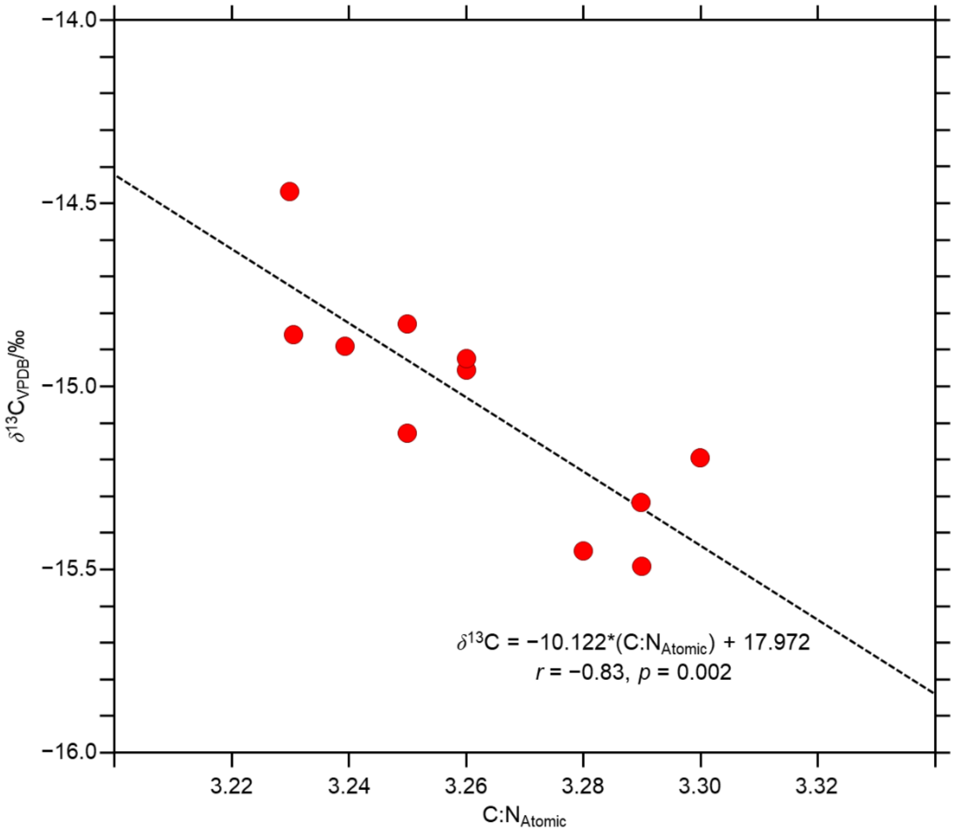


**Figure S1**. Relationship between the $\delta^{13}$C and C:N$_{Atomic}$ of modern polar bear bone collagen analyzed in this study.

The $\delta^{13}$C values of the modern polar bear bone collagen were corrected to account for lipid contamination according to the equation of a line in Figure S1 as described in the main text.

**Results**

The stable isotope and elemental compositions for the ancient polar bears analyzed in this study are presented in Table S2. The stable isotope and elemental compositions for the modern polar bears analyzed in this study are presented in Table S3.
