## Supplement S2 Archaeological Data for "Unprecedented shift in Canadian High Arctic polar bear food web unsettles four millennia of stability"

| Sample ID | Cultural Affiliation | Site | ~ Site Age (cal yrs BP) | Location | $\delta^{13}C_{VPDB}$/‰ | $\delta^{15}N_{AIR}$/‰ | wt %C | wt %N | $C:N_{Atomic}$ |
|---|---|---|---|---|---|---|---|---|---|
| 6004 | Pre-Dorset | QkHn-13 | 4150-3645 | Devon Island | −12.79 | +20.78 | 44.7 | 16.4 | 3.18 |
| 6025 | Pre-Dorset | QkHn-13 | 4150-3645 | Devon Island | −13.13 | +22.06 | 45.4 | 16.6 | 3.19 |
| 6087 | Pre-Dorset | QkHn-13 | 4150-3645 | Devon Island | −13.37 | +23.54 | 45.4 | 16.6 | 3.19 |
| 6114 | Pre-Dorset | QkHn-13 | 4150-3645 | Devon Island | −13.18 | +22.09 | 44.8 | 16.3 | 3.21 |
| 6177 | Pre-Dorset | RcJu-1 | 3565-3405 | Devon Island | −13.15 | +21.60 | 44.0 | 15.9 | 3.23 |
| 6180 | Pre-Dorset | RcJu-1 | 3565-3405 | Devon Island | −13.82 | +23.91 | 45.3 | 16.4 | 3.22 |
| 6200 | Pre-Dorset | RbJu-1 | 4285-3935 | Devon Island | −13.76 | +23.35 | 45.6 | 16.6 | 3.20 |
| 6211 | Pre-Dorset | RbJu-1 | 4285-3935 | Devon Island | −13.13 | +21.99 | 45.6 | 16.6 | 3.20 |
| 6750 | Pre-Dorset | RbJu-1 | 4285-3935 | Devon Island | −14.04 | +23.24 | 45.5 | 16.3 | 3.26 |
| 6752 | Pre-Dorset | RbJu-1 | 4285-3935 | Devon Island | −13.19 | +22.21 | 43.9 | 15.9 | 3.22 |
| 4507 | Late Dorset | QjJx-10 | 1545-550 | Little Cornwallis Island | −14.01 | +23.35 | 45.4 | 16.6 | 3.19 |
| 4536 | Late Dorset | QjJx-10 | 1545-551 | Little Cornwallis Island | −14.02 | +23.58 | 45.0 | 16.4 | 3.20 |
| 4550 | Late Dorset | QjJx-10 | 1545-552 | Little Cornwallis Island | −14.39 | +23.24 | 44.9 | 15.9 | 3.29 |
| 4557 | Late Dorset | QjJx-10 | 1545-553 | Little Cornwallis Island | −13.47 | +23.81 | 44.9 | 16.4 | 3.19 |
| 4617 | Late Dorset | QjJx-10 | 1545-554 | Little Cornwallis Island | −13.45 | +23.10 | 45.6 | 16.4 | 3.24 |
| 4653 | Late Dorset | QiLa-3 | 935-725 | Little Cornwallis Island | −13.76 | +20.77 | 45.2 | 16.2 | 3.25 |
| 4655 | Late Dorset | QiLa-3 | 935-725 | Little Cornwallis Island | −13.60 | +20.09 | 46.2 | 16.8 | 3.21 |
| 4676 | Late Dorset | QiLa-3 | 935-725 | Little Cornwallis Island | −13.37 | +20.78 | 44.9 | 16.3 | 3.21 |
| 4682 | Late Dorset | QiLa-3 | 935-725 | Little Cornwallis Island | −13.65 | +21.04 | 43.2 | 15.1 | 3.34 |
| 4702 | Late Dorset | QjJx-1 | 1045-545 | Little Cornwallis Island | −12.96 | +21.68 | 44.9 | 16.3 | 3.20 |
| 4717 | Late Dorset | QjJx-1 | 1045-545 | Little Cornwallis Island | −13.58 | +22.71 | 45.6 | 16.4 | 3.24 |
| 4755 | Late Dorset | QjJx-1 | 1045-545 | Little Cornwallis Island | −13.03 | +21.91 | 45.3 | 16.5 | 3.20 |
| 4766 | Late Dorset | QjJx-1 | 1045-545 | Little Cornwallis Island | −12.84 | +21.01 | 45.9 | 16.6 | 3.22 |
| 5827 | Late Dorset | QjLd-17 | 900-675 | Little Cornwallis Island | −13.72 | +23.64 | 43.1 | 15.6 | 3.22 |
| 5852 | Late Dorset | QjLd-17 | 900-675 | Little Cornwallis Island | −13.57 | +23.56 | 45.7 | 16.8 | 3.17 |
| 6594 | Thule | RbJr-1 | 675-500 | Devon Island | −13.50 | +22.89 | 45.0 | 16.3 | 3.22 |
| 6620 | Thule | RbJr-1 | 675-500 | Devon Island | −13.58 | +22.03 | 45.6 | 16.5 | 3.22 |
| 6686 | Thule | RbJr-1 | 675-500 | Devon Island | −13.77 | +21.51 | 45.6 | 16.4 | 3.24 |
| 8109 | Thule | PcJq-5 | 925-510 | Somerset Island | −13.71 | +20.51 | 45.4 | 15.8 | 3.35 |
| 8120 | Thule | PcJq-5 | 925-510 | Somerset Island | −13.40 | +21.50 | 45.2 | 16.4 | 3.21 |

| | | | | | | | | | |
|---|---|---|---|---|---|---|---|---|---|
| 8345 | Thule | PcJq-5 | 925-510 | Somerset Island | −13.35 | +22.55 | 45.4 | 16.4 | 3.23 |
| 8539 | Thule | PcJq-5 | 925-510 | Somerset Island | −12.62 | +21.95 | 40.2 | 14.6 | 3.22 |
| 8500 | Thule | PeJr-1 | 640-505 | Somerset Island | −13.29 | +22.73 | 45.2 | 16.1 | 3.27 |
| 8517 | Thule | PeJr-1 | 640-505 | Somerset Island | −12.80 | +22.66 | 40.0 | 14.5 | 3.22 |
| 8529 | Thule | PeJr-1 | 640-505 | Somerset Island | −12.94 | +21.93 | 44.0 | 15.9 | 3.23 |
