## Supplement S3 Modern Data for "Unprecedented shift in Canadian High Arctic polar bear food web unsettles four millennia of stability"

| Sample ID | Collection Year | $\delta^{13}C_{VPDB}$/‰ | $\delta^{13}C_{lipid\ corrected\ VPDB}$/‰[1] | $\delta^{13}C_{corr\ VPDB}$/‰[2] | $\delta^{15}N_{AIR}$/‰ | wt %C | wt %N | $C:N_{Atomic}$ |
|---|---|---|---|---|---|---|---|---|
| 220 | 1998 | −14.86 | −14.86 | −14.10 | +22.22 | 40.1 | 14.5 | 3.23 |
| 221 | 1998 | −14.89 | −14.79 | −14.13 | +22.20 | 40.8 | 14.7 | 3.24 |
| 224 | 1998 | −15.19 | −14.48 | −14.43 | +21.94 | 40.5 | 14.3 | 3.30 |
| 231 | 1999 | −14.47 | −14.47 | −13.69 | +21.91 | 40.2 | 14.5 | 3.23 |
| 232 | 1999 | −14.83 | −14.62 | −14.05 | +20.25 | 39.9 | 14.3 | 3.25 |
| 233 | 1999 | −14.95 | −14.65 | −14.17 | +21.82 | 38.9 | 13.9 | 3.26 |
| 234 | 1999 | −15.45 | −14.94 | −14.67 | +23.02 | 39.9 | 14.2 | 3.28 |
| 226 | 2001 | −15.31 | −14.70 | −14.48 | +21.80 | 40.3 | 14.3 | 3.29 |
| 227 | 2001 | −14.93 | −14.63 | −14.10 | +22.17 | 40.5 | 14.5 | 3.26 |
| 219 | 2002 | −15.49 | −14.88 | −14.64 | +21.35 | 39.5 | 14.0 | 3.29 |
| 237 | 2007 | −15.13 | −14.93 | −14.16 | +21.75 | 42.9 | 15.4 | 3.25 |

*1.* $\delta^{13}C$ values corrected for the presence of lipid contaminants as described in the main text.

*2.* $\delta^{13}C$ values corrected for the reduction in oceanic $CO_2$ $\delta^{13}C$ values due to industrialization as described in the text. These $\delta^{13}C$ values have also been corrected for the presence of lipid contaminants.
